# Use of RetroNectin in studies requiring in vitro HIV-1 infection of human hematopoietic stem/progenitor cells

**DOI:** 10.1101/142075

**Authors:** Tetsuo Tsukamoto, Seiji Okada

## Abstract

Human immunodeficiency virus (HIV) causes damage, directly or indirectly, to the whole hematopoietic system including CD34^+^ hematopoietic stem/progenitor cells (HSPC). CXCR4-tropic strains of HIV-1 may be potent to affect the function of CD34^+^CXCR4^+^ progenitor cells either by infecting the cells or by modifying the dynamics of more differentiated hematopoietic cells. However, CD34^+^ cells are known for the resistance to HIV-1 infection in vitro, restricting the detailed analysis of the impact of HIV upon HSPC. Here the authors report a use of RetroNectin, a recombinant fibronectin fragment used for gene transfer with lentiviral vectors, to overcome the limitation.

---

The chemokine receptor CXCR4 is a co-receptor of human immunodeficiency virus (HIV) (Feng et al., 1996). Physiologically, CXCR4 interacts with stromal cell-derived factor 1 (SDF-1) and allows CXCR4-expressing cells to home to the loci where SDF-1 is highly expressed (Kucia et al., 2004). For example, the roles of SDF-1 and CXCR4 are essential in human stem cell homing and repopulation of the host with differentiated hematopoietic cells (Kucia et al., 2005; Lapidot and Kollet, 2002). SDF-1 is also produced by thymus epithelial cells and plays an important role in migration of immature progenitors in the thymus (Plotkin et al., 2003). In contrast to another HIV-1 co-receptor CCR5, which has a widely-distributed deletion mutant called Δ 32 in Caucasian populations (Stephens et al., 1998), the physiological roles of CXCR4 are indispensable.

Therefore, it will be important to better understand the impact of CXCR4-tropic HIV-1 infection upon hematopoiesis-related events including T cell development in the thymus. Bone marrow abnormalities such as dysplasia and abnormal hematopoietic cell development are frequently observed in HIV infected individuals (Tripathi et al., 2005). Thymus dysfunction occurs during HIV disease and its rapid progression in infants is associated with prenatal HIV infection (Ye, Kirschner, and Kourtis, 2004).

HIV-1 has recently been shown to infect human CD34+ hematopoietic progenitor cells (Carter et al., 2011; Carter et al., 2010). On the other hand, hematopoietic stem/progenitor cells (HSPC) limit HIV infection in different ways. One is the extremely low expression levels of the receptors on these cells (Carter et al., 2011). Besides, some reports indicate mechanisms restricting HIV-1 prior to integration (Griffin and Goff, 2015). These may prevent researchers from detailed in vitro analysis of CD34+ cells in the presence of HIV-1. To overcome the limitations, the authors propose a method to mediate HIV-1 entry to CD34^+^ cells by using RetroNectin (RN), a recombinant fibronectin fragment that enhances retroviral-mediated gene transduction by aiding the co-localization of target cells and virions.

RetroNectin has been tested with vesicular stomatitis virus glycoprotein (VSV-G), murine ecotropic/amphotroic/xenotropic retrovirus envelopes (Lee, Sadelain, and Brentjens, 2009; Relander et al., 2001; Sakuma et al., 2010), and gibbon ape leukemia virus (GaLV) envelopes (Chono et al., 2001). However, RN’s ability to help CXCR4-tropic HIV-1 envelopes at the viral entry to target cells has not been focused. The authors demonstrate that cord-derived CD34^+^ cells can be stably HIV-infected for weeks.

Umbilical cord blood (UCB) samples were collected at Fukuda Hospital, Kumamoto, Japan following informed consent. Cord blood mononuclear cells were isolated and CD34^+^ cells were selected using human CD34 microbeads and LS columns (Miltenyi Biotec, Tokyo, Japan). The purity was always more than 92 % on flow cytometry. The purified CD34^+^ cells (2×10^5^) were seeded in a non-tissue culture-treated Falcon 48-well plate (catalog #351178, Corning, Tokyo, Japan) coated with RetroNectin (Takara Bio Inc., Tokyo, Japan) at a concentration of 10 μg/mL following the manufacturer’s protocol. The media for infection and culture was MEM-α supplemented with 10 % heat inactivated fetal bovine serum (FBS), 5 ng/mL of recombinant human interleukin 7 (IL-7), and 5 ng/mL of FMS-like tyrosine kinase 3 ligand (Flt3L). After adding 200 ng (p24) of HIV-1_NL4–3_ (Adachi et al., 1986), the plate was kept at 34 °C, centrifuged at 1,200 g for 30 minutes, and cultured overnight at 37 °C. Cells were then transferred to a coculture with a mouse stromal cell line OP9-DL1 (Fig 1A) (Zuniga-Pflucker, 2009).

**Figure 1.**
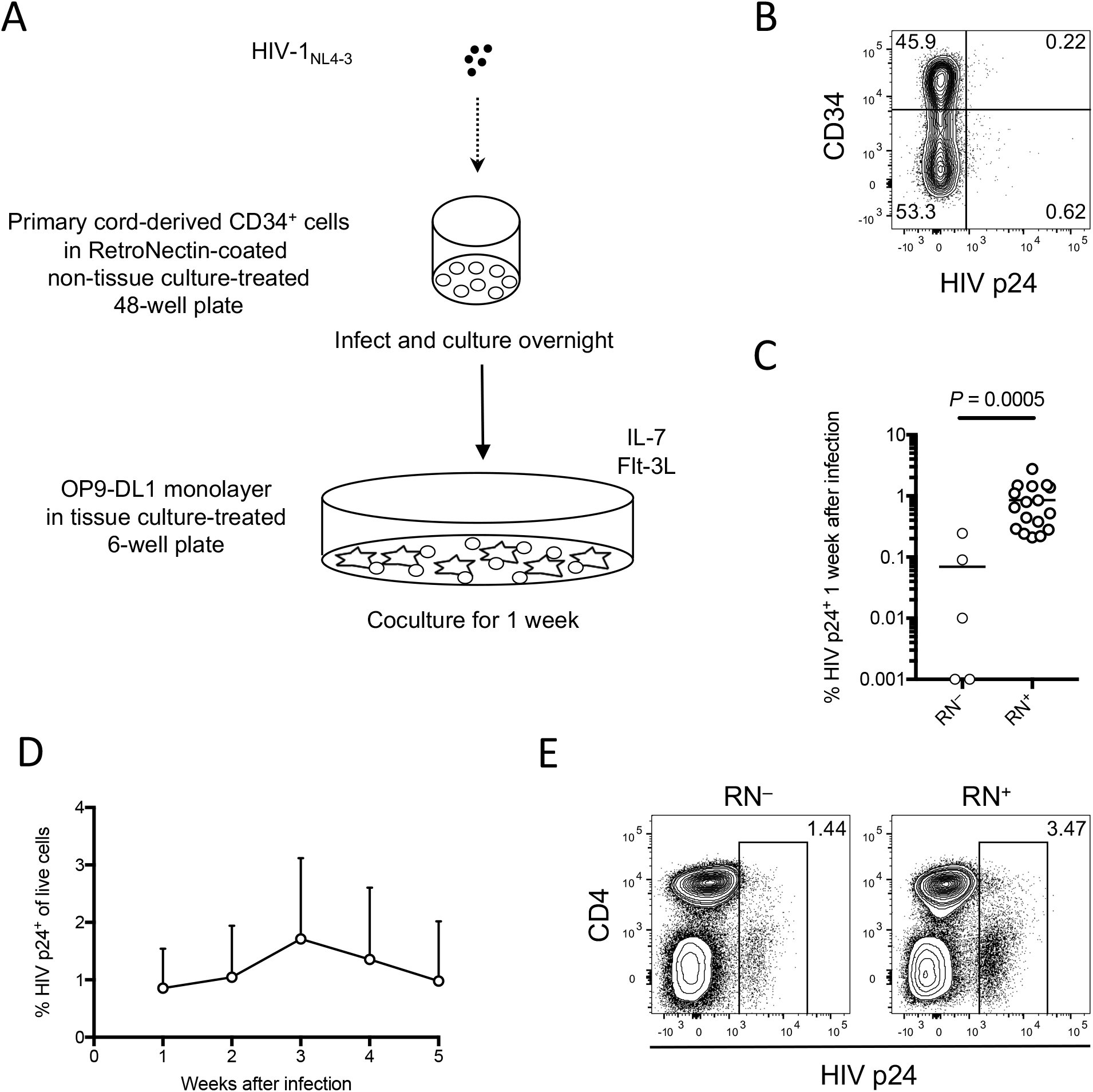
RetroNectin (RN)-assisted infection of primary cord-derived CD34^+^ cells with a CXCR4-tropic strain of HIV-1. (A) Schematic representation of the procedure. CD34^+^ cells were infected with HIV-1_NL4–3_ in a RN-coated plate, and cocultured with OP9-DL1 for a week. (B) Intracellular HIV-1 p24 expression was measured a week after infection. HIV-1 p24^+^ cells were found in both CD34^+^ and CD34^−^ fractions of HIV-infected samples. A representative plot is shown. (C) RN-assisted CD34^+^ cells (n=17) showed much higher HIV-1 p24+ frequencies than RN-unassisted CD34^+^ cells (n=5) at week 1 post infection. The statistical analysis was done by a Mann-Whitney test. (D) Intracellular HIV-1 p24 expression levels in RN-assisted HIV-infected CD34^+^ cells were followed for 5 weeks (n=17). (E) Cord blood mononuclear cells were resuspended in RPMI1640 media supplemented with 10 % FBS and 20 IU/mL recombinant human IL-2. Cells were infected with HIV-1_NL4–3_ in a 48-well plate with or without RN coating, cultured for a week, and tested for intracellular HIV-1 p24 expression on flow cytometry. The HIV-1 p24^+^ frequency of RN-assisted sample (right panel) was compared to that of RN-untreated sample (left panel).

Samples were analyzed at week 1 post infection and coculture. Cells were stained with anti-human CD34 APC (BD Biosciences, Tokyo, Japan) and anti-HIV-1 p24 PE (Beckman Coulter, Tokyo, Japan), and analyzed on flow cytometry (FACS LSR II, BD Biosciences). Intracellular HIV-1 p24^+^ frequencies were calculated by subtracting the background staining levels without HIV-1 infection. Both CD34^+^ and CD34^−^ cells were detected in the flow cytometric analysis, suggesting that cord-derived CD34^+^ cells partly underwent in vitro differentiation in the coculture (Fig 1B). HIV-1 p24^+^ cells were found in both CD34^+^ and CD34^−^ fractions, showing that HIV-1 successfully infected CD34^+^ cells. Next, comparison of HIV-1 p24+ frequencies at week 1 post infection was done between samples using and not using RN for infection. The average HIV-1 p24^+^ frequencies in RN-treated samples were close to 1 %, while HIV-1 infection without RN coating didn’t always result in productive infection of CD34^+^ cells (Fig 1C). In all the RN-treated samples tested (n=17), the intracellular p24+ frequencies were higher than 0.20 (Table 1), manifesting the reliability of the method for infection of relatively virus-resistant CD34^+^ cells. With all the samples tested, RN-assisted HIV-1 infection of CD34^+^ cells lasted at least for 5 weeks (Fig 1D). It was also checked that merely treating CD34^+^ cells with RN without HIV-1 infection didn’t cause any changes to the cells in terms of the whole cell counts and CD34/CXCR4 expression levels, when compared to RN-untreated cells (data not shown).

**Table 1.**
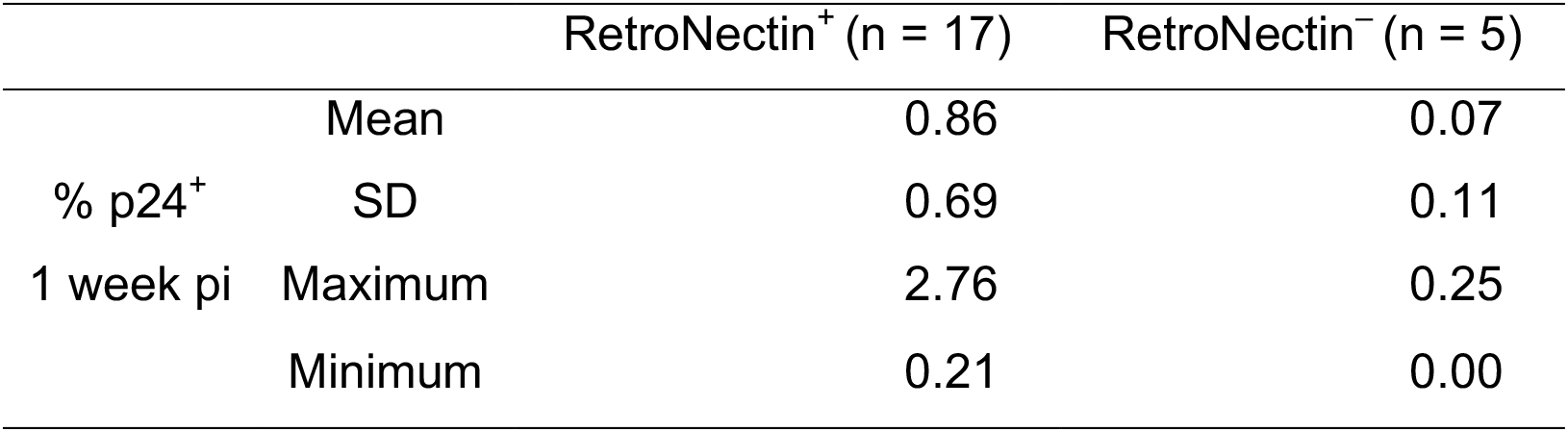
Summary for the HIV infection of CD34^+^ cells with or without RN coating. SD: standard deviation. pi: post infection.

To confirm the enhancing effect of RN on HIV-1 entry, the whole cord blood mononuclear cells were infected with HIV-1_NL4–3_ in an RN-coated or -uncoated well. After the centrifugation as above, followed by a culture for a week in RPMI1640 media supplemented with 10 % FBS and 20 IU/mL IL-2, cells were stained with anti-human CD4 PerCP-Cy5.5 (BioLegend, Tokyo, Japan) and anti-HIV-1 p24 PE, and analyzed on flow cytometry. As shown in Fig 1E, RN also supported HIV-1 infection of blood mononuclear cells. However, the intracellular HIV-1 p24^+^ rate without RN (1.44 %) was much higher than the mean HIV^+^ rate in CD34^+^ cells infected without RN (0.07 %, Table 1), meaning that RN coating might be also of some help in HIV infection of blood mononuclear cells, although not essential. A possible role for RN to activate T cells should also be noted (Ishikawa et al., 2014; Li et al., 2015).

Some might see the finding as a matter of course, since RN coated plate readily helps lentiviral transduction of CD34^+^ cells. However, lentiviral vectors are usually pseudotyped with VSV-G that works as a strong drive for viral entry to the cells. In the current method, on the other hand, HIV-1 uses its native receptors CD4 and CXCR4. Their expression levels are extremely limited on cord-derived CD34^+^ cells. In addition, the experiment here didn’t include strong cytokine stimulation required for lentiviral transduction of HSPC (Uchida et al., 2011). Even in this condition, as shown in the results, RN is powerful enough to allow stable HIV infection of human cord-derived CD34^+^ cells.

The present study didn’t test a CCR5-tropic strain of HIV-1. CCR5-tropic immunodeficiency viruses cause massive depletion of memory CD4^+^ T cells in the acute phase and are deeply linked to AIDS development (Mattapallil et al., 2005; Okoye and Picker, 2013). Even if those strains seem to have limited direct effect on HSPC, it might be worth testing whether they can infect HSPC efficiently in vitro. Similarly, it will be exciting to apply the current method to other retroviruses, including simian immunodeficiency viruses and murine leukemia viruses, as well as other viruses that can utilize RN for entering their inefficient target cells.

The simple and easy method of HIV infection using RN is highly effective on infection of HIV-resistant cells. With primary human cord-derived CD34^+^ cells, the RN coating of a plate enhanced the average HIV+ rate at week 1 by more than 10 folds. All the samples tested with RN-coated plate showed stable HIV-1 infection for 5 weeks. The method may be applicable to any combination of virus-resistant suspension cells and a virus like retroviruses, bringing long-term in vitro analyses of infected cells, including rare or precious cell samples, close to researchers’ hands.

## Acknowledgements

We cordially thank Drs. Kazuo Matsui and Shoichi Kawakami of Fukuda Hospital, Kumamoto, Japan for their help on cord blood sampling. This work was partly supported by grants from the following organizations: Japan Agency for Medical Research and Development (Research program on HIV/AIDS, No. 16fk0410108h0001); Ministry of Education, Culture, Sports, Science and Technology, Japan (Grants-in-Aid for Science Research, No. 25114711); National Health and Medical Research Council, Australia (Project grant). Special thanks to Prof. Anthony D. Kelleher and Dr. Kazuo Suzuki of the Kirby Institute for infection and immunity for society, UNSW Australia for their support for the whole project, which was planned by UNSW Australia and hosted by Kumamoto University, Japan, from distance.

